# Two Novel Rod-Like Circular RNAs from the Oroarctic Tundra Soil Display Candidate Ribozyme-Like Features: An In Silico Characterization

**DOI:** 10.64898/2026.07.27.740603

**Authors:** Noah Williams, Garrett Wolfram, Alice Zhang, Maya Tessler, Geunhwa Jung

## Abstract

The recent discovery of obelisks, novel deltaviruses, and other viroid-like circular RNAs has greatly expanded the known diversity of autonomous RNA elements. However, cold terrestrial ecosystems remain comparatively undersampled for these molecules. Using a structure-based modification of the Tormentor pipeline, we assembled two putative non-coding rod-like circular RNAs from winter metatranscriptomes of oroarctic tundra soil collected near Kilpisjärvi, Finland. Sequence-based searches identified no meaningful homologs for either RNA. Structure-based analyses identified affinities to deltavirus-like ribozymes in Rod 1 and to hammerhead ribozymes in Rod 2, with structural compatibility confirmed by integration into an established covariation model. Rod 1 was additionally detected in independent alpine metatranscriptomic datasets, whereas Rod 2 was not identified beyond the original sample. Collectively, these results identify Rod 1 as a strong candidate viroid-like RNA bearing a putative ribozyme-like structure motif that warrants experimental validation, while Rod 1 remains a preliminary candidate requiring further confirmation.

## Introduction

The diversity of viroid-like and other circular RNA has expanded rapidly with the discovery of the broader deltavirus lineage, obelisks, and numerous previously unrecognized circular RNA elements across diverse environments (Denis et al., 2026; Raza et al., 2025; López-Simón et al., 2025). These discoveries have demonstrated that many biologically relevant RNA elements cannot be identified through sequence homology alone, motivating the development of structure-based discovery approaches. Low temperature environments are of particular interest because RNA folding, stability, and catalytic activity are strongly influenced by temperature (Attwater et al., 2013). Despite their importance for understanding of RNA evolution and adaptation, arctic and alpine microbial ecosystems remain relatively poorly characterized due to logistical changes associated with sampling and a historical environment of agricultural, medical, and industrial relevance (Herndon et al., 2015).

Obelisks were initially identified in the human microbiome and have since been shown to occur widely in oceanic and other environmental samples (Zheludev et al., 2024; López-Simón et al., 2025). More recently, a novel viroid-like RNA associated with a solanaceous crop, was discovered using a search-based strategy rather than conventional sequence homology, highlighting the power of structural approaches for identifying highly divergent RNA elements (Zheludev et al., 2024; Kremer & De Barros., 2024; Raza et al., 2025). Although most subsequent searches focused on crop-associated datasets, a limited number of environmental metatranscriptomes were also examined.

Here, we report the computational characterization of two previously undescribed rod-like circular RNAs identified in a winter metatranscriptome from oroarctic tundra soil collected near Kilpisjärvi, Finland. We evaluated these RNAs using sequence homology searches, structural comparisons, covariance analyses, and distribution across independent metatranscriptomatic databases. Our results provide evidence that one element, Rod 1, possess structural features consistent with known ribozyme families and is detected in geographically independent alpine datasets, whereas Rod 2 remains a more tentative candidate pending additional evidence.

## Methods

### Data Source & Assembly

The program *VNOM* as part of the *Tormentor* pipeline was used to de novo assemble double paired RNA-seq data from the SRA database (Zheludev et al., 2024 ; Kremer & De Barros., 2024). Criteria were set to 70% circularity and a minimum read depth of 30, however the final step of the pipeline, which searches for homology to the protein Oblin1, was bypassed to target all highly self-pairing RNAs. Resulting structures from RNAfold that passed the 70% self-pairing criteria were visually assessed to screen for rod-like morphology. Structures were excluded if any branching stem exceeded one-third of the total structure length. (Lorenz et al., 2011). YASS was used to evaluate sequence repetitiveness to screen out sequencing artifacts (Noé & Kucherov, 2005). Hundreds of datasets were selected for relevance to plant pathology alongside a known obelisk positive sample SRR5949245. An ad hoc selection of environmental samples was opportunistically included (n<20). Among the environmental samples was SRX28877771. The sample was collected in Kilpisjärvi, Finland as part of a study of microbial community composition in the oroarctic tundra as described in Sirja et al. (2025).

### Study site and conditions

The sample was collected from winter soil near Kilpisjärvi, Finland, a high-elevation tundra environment, from a seasonally dry wetland by Sirja et al., (2025). The soil layer is shallow (<50 cm) and organic rich. pH was 5.05, soil organic matter content was 81.45%, and C:N ratio was 14.61 (SOM and C:N measured in summer). Soil temperature was -1.5C and snow depth was 62 cm. Samples were taken at ∼5cm soil depth.

### Homology Check

A 1003bp rod-like structure (MFE of -315.40 kcal/mol and a GC 48.65%) (Figure 1) and A 1063bp rod-like structure (MFE **-**362.20 kcal/mol and a GC 51.65%) (Figure 2) were found and characterized further. Both rods were checked for potential ORF frames using the NCBI ORF finder. YASS was used as a first search for repetitions in both rods. Sequences were then searched through *Censor* against a peer reviewed database of transposons (Kohany et al., 2006).*Uniprot* was used to search for potential protein matches (*Oblin* 1 and *Oblin* 2 are within the Uniprot database). NCBI MegaBLAST was performed against the core nucleotide database and the four experimental databases. Paired discontinuous MegaBLAST searches were performed against a representative Epsilon virus (OV121072.9), Zeta virus (OV121037.13), hop latent viroid (X07397.1), and the hepatitis D deltavirus (NC_076103.1), as well as for Rod 1 and Rod 2 to each other.

**Figure 1.**
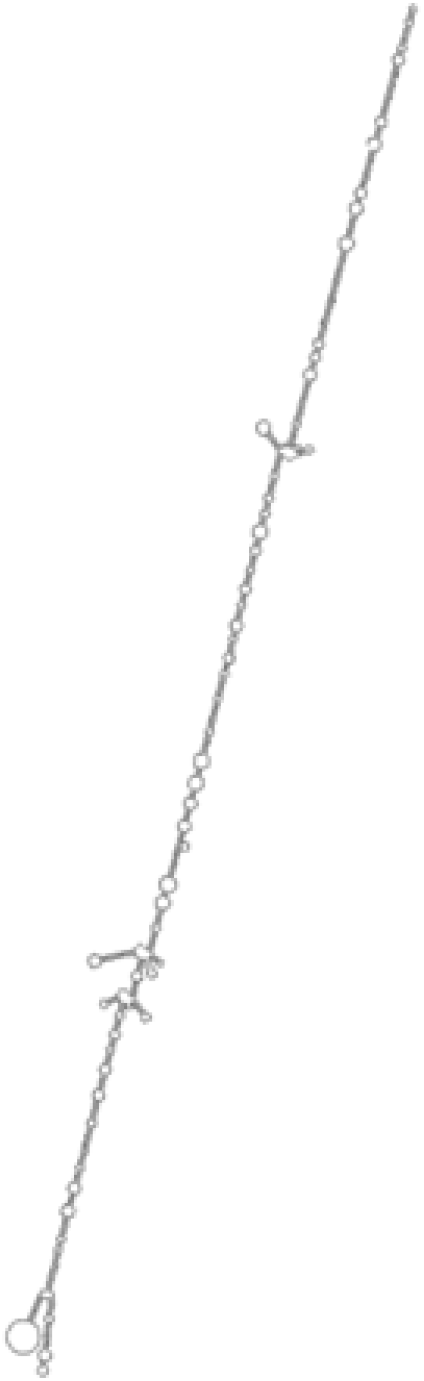
Minimum free energy (MFE) structure of Rod 1 predicted by ViennaRNA RNAfold (ΔG = -315.40 kcal/mol).

**Figure 2.**
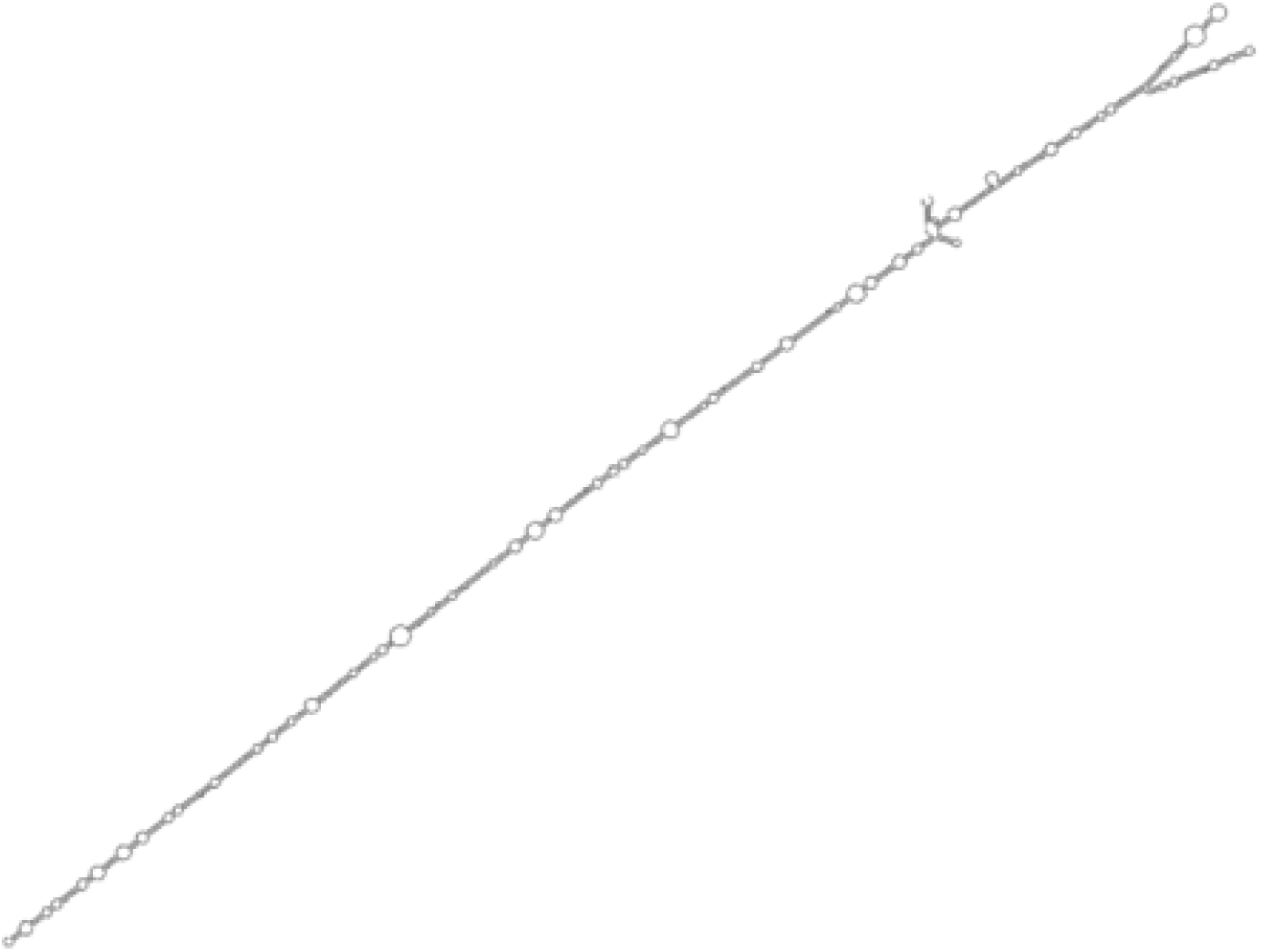
Minimum free energy (MFE) structure of Rod 2 predicted by ViennaRNA RNAfold (ΔG = -362.20 kcal/mol).

### Structural Search and Analysis

Since sequence homology had no results Infernal was used for structure based searches against RFAM families (Nawrocki et al., 2009). The regions surrounding the two significant hits from Rod 1 (100bp, 86bp) and the notable hit from Rod 2 (66bp) were run through the RNAhub pipeline ending in R-scape (Rivas et al., 2020).

The three regions surrounding the hits were input to *RNAnneal*, a biophysics based modeller (Herron et al.,2026) and assessed for fold patterns. IPKNOT using the Boltzmann model and pKiss were used to validate the results, with pKiss reported as the more accurate tool for pseudoknot prediction (Miklós et al., 2005;Theis et al., 2010; Gong et al., 2024). All tests are by default based on 37 degree temperatures outside the range of temperatures the origin sample existed in. To correct for this pKiss was used to predict pseudoknots at temperature ranges actually feasible ranging from -2C to 20C, with the 37C prediction retained for reference. If a conformation change was observed between 20C and 37C the tested range would be expanded. To validate the conformational changes pKiss mode “probs” was used to look at non-optimal structures.

All three sequences were run through all Pebblescout and Metagraph datasets, two kmer based search tools for assembled SRA data (Edgar et al., 2022; Shiryev et al., 2024). A MegaBLAST of the sequence was run against the SRA datasets with less than 100% similarity (from the pebblescout search) in search of reads with divergent basepairs. The other SRA datasets from the original project were searched via MegaBLAST. Rod 1 region 1 and Rod 2 region 1 were searched against the 20 SRA datasets in (SRP653358), a study too recent to have been assembled in pebblescout, metatranscriptomics data from Alaskan arctic soil.

## Results

Thirteen highly self pairing (>70%) RNAs were assembled from the SRX28877771 RNA-seq data, of which two met the criteria for a rod-like structure. The obelisk positive sample SRR5949245 assembled one highly self pairing RNA which also met the criteria for a rod-like structure, confirming the pipeline was functioning. For Rod 1 no open reading frames were predicted that were longer than 78 amino acids and the sequence is putatively non-coding. For Rod 2 the longest predicted open reading frame was 94 amino acids long and is also putatively non-coding. In Rod 1, a self-alignment was identified between positions 22–195 and 727–895 (E = 2.7×10^−4^, bit-score 31.8), oriented in reverse complement and consistent with a diverged inverted repeat of roughly 170 nt, otherwise the sequence was non-repetitive. In Rod 2 three self alignments were identified, however all had E values greater than 1, 320-382 and 774-836 (E value 1.06, bit-score = 20.02), 798-826 and 324-349 (E value 5.25, bit-score = 17.72), and 615-737 and 8-130 (E value 6.26, bit-score 17.46). The Censor search revealed for Rod 1 a short 56bp overlap with the non ltr-transposon Ingi-1_LMi, and no hits for Rod 2 (Bao & Jurka., 2014). The UniProt search for protein matches also had no significant hits for both sequences. The MegaBLAST searches revealed for Rod 1 marginal hits to genome assemblies of *Phosphuga atrata* 34/34bp OX155901.1 and *Laparocerus anagae* 36/37bp OZ344904.1, no hits for Rod 2. Paired discontinuous MegaBLAST searches all found no alignment.

Infernal had two significant hits for Rod 1. Hits were from the ribozymes of chimpanzee deltavirus like ribozyme 7qr3C Positions 77 to 159 (E-value 0.0056) and from hepatitis deltavirus ribozyme 3nkb8 Positions 878 to 925 (E-value 0.0044) which both overlap with the self-alignment identified by YASS (Przytula-Mally et al.,2022). For Rod 2 one hit was notable but non-significant to the hammerhead ribozyme 1hmhA E positions 502-535 (E-value 0.078). The RNAhub pipeline revealed that Rod 1 region 1 matched the CPEB3 ribozyme with an E-value of 0.00017. The modelled family with the Rod 1 region 1 sequence had significant covariation, within the family model into which it was incorporated at sites 131/145 p = .0006 and 133/145 p = .0017 (Figure 3). For Rod 1 region 2 no significant matches were found, as a result no alignment could be built and a configuration was forced in R-scape (Figure 4). For Rod 2 region 1 the RNAhub pipeline found a significant hit E value 6.1e-07 to RF00008 Hammerhead ribozyme (type III) with ten significant covarying basepairs when incorporated into the family model (Figure 5).

**Figure 3.**
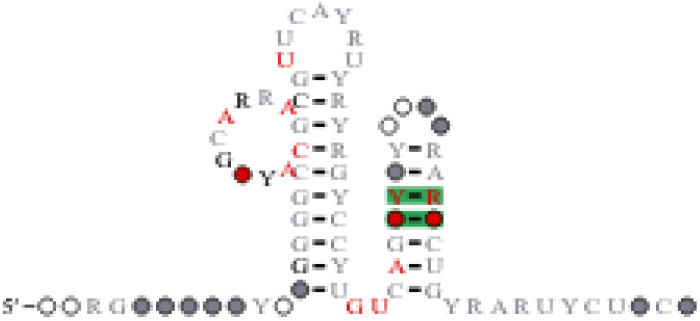
R-scape covariation analysis of Rod 1 region 1 aligned to the Rfam CPEB3 ribozyme family. Significantly covarying base pairs are highlighted in green.

**Figure 4.**
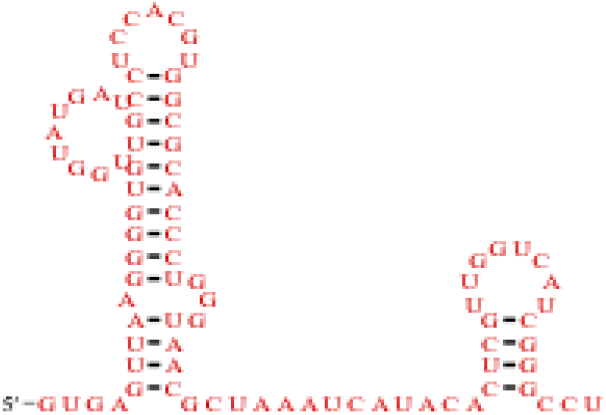
R-scape analysis of Rod 1 region 2 using a manually constrained alignment. No significant match to any known Rfam structural family was identified.

**Figure 5.**
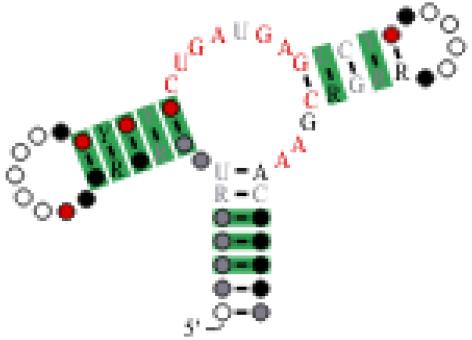
R-scape covariation analyses of Rod 2 region 1 aligned to the Rfam RF00008 hammerhead ribozyme (type III) family. Significantly covarying base pairs are highlighted in green.

The RNAAnneal predicted structure for Rod 1 region 1 had a pair of pseudoknots in the region, supported across five simulations, while Rod 1 region 2 lacked any apparent pseudoknots. Rod 2 region 1 had 0 pseudoknots predicted across five simulations. IPKNOT using the Boltzmann model predicted for Rod 1 region 1 one pseudoknot, and for Rod 1 region 2 three pseudoknots (one nested pair). For Rod 2 region 1 no pseudoknots were predicted by the Boltzmann model The pKiss mfe model at default settings predicted two pseudoknots for Rod 1 region 1, a nested pair for Rod 1 region 2, and 3 pseudoknots for Rod 2 region 1. Testing the pKiss mfe model across biologically feasible temperatures conformational shifts were observed in Rod 1 region 1 at or below 5C, and for Rod 1 region 2 below 19C, increasing predicted pseudoknot count or shifting pseudoknot location respectively. However, when looking at the probability of non-optimal structures, a gradual shift at 20C is observed for both, with Rod 1 region 1 showing a clear loss of pseudoknot 3 with rising temperatures and Rod 1 region 2 showing a shift in structure at high and low temperatures (Figure 6 & Figure 7). For Rod 2 region 1 temperatures ranging from -2C to 20C were explored, all predicted no pseudoknots while the 37 degree model predicted 3 pseudoknots. To determine the conformational shift point the temperature range was expanded to 30C, and a shift was observed at 24C and higher. However, when looking at non-optimal structures pseudoknot presence is consistent from -2C onwards but several positional differences are competing, with one 3 pseudoknot configuration dominating with 95% probability at 37C. The structure seems to only stabilize at temperatures outside the biologically relevant range.

**Figure 6.**
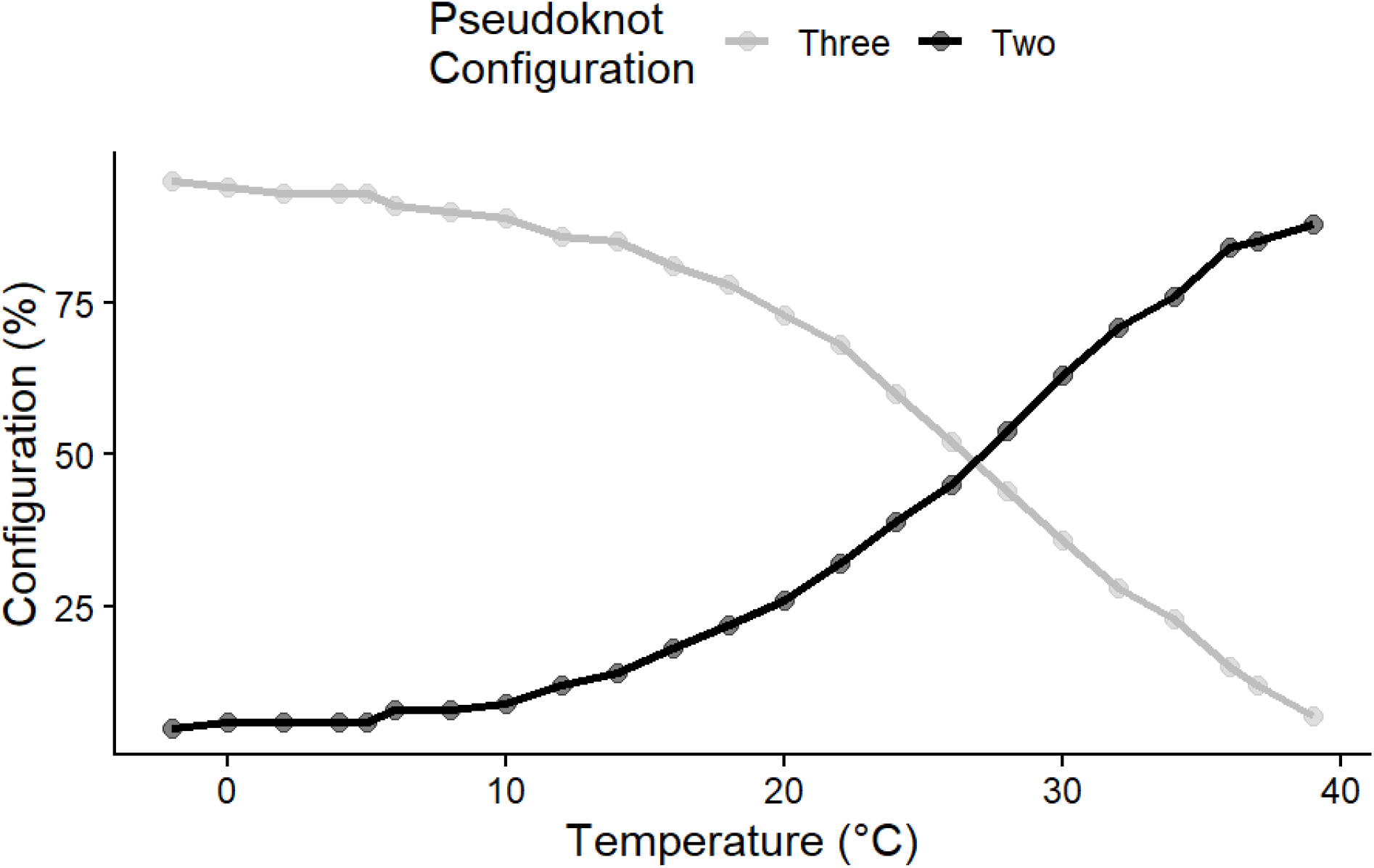
Predicted pseudoknot configuration of Rod 1 region 1 generated using the pKiss “probs” model across temperatures ranging from -2C to 39C. Configurations were grouped according to the presence or absence of the third pseudoknot at the 3’ end.

**Figure 7.**
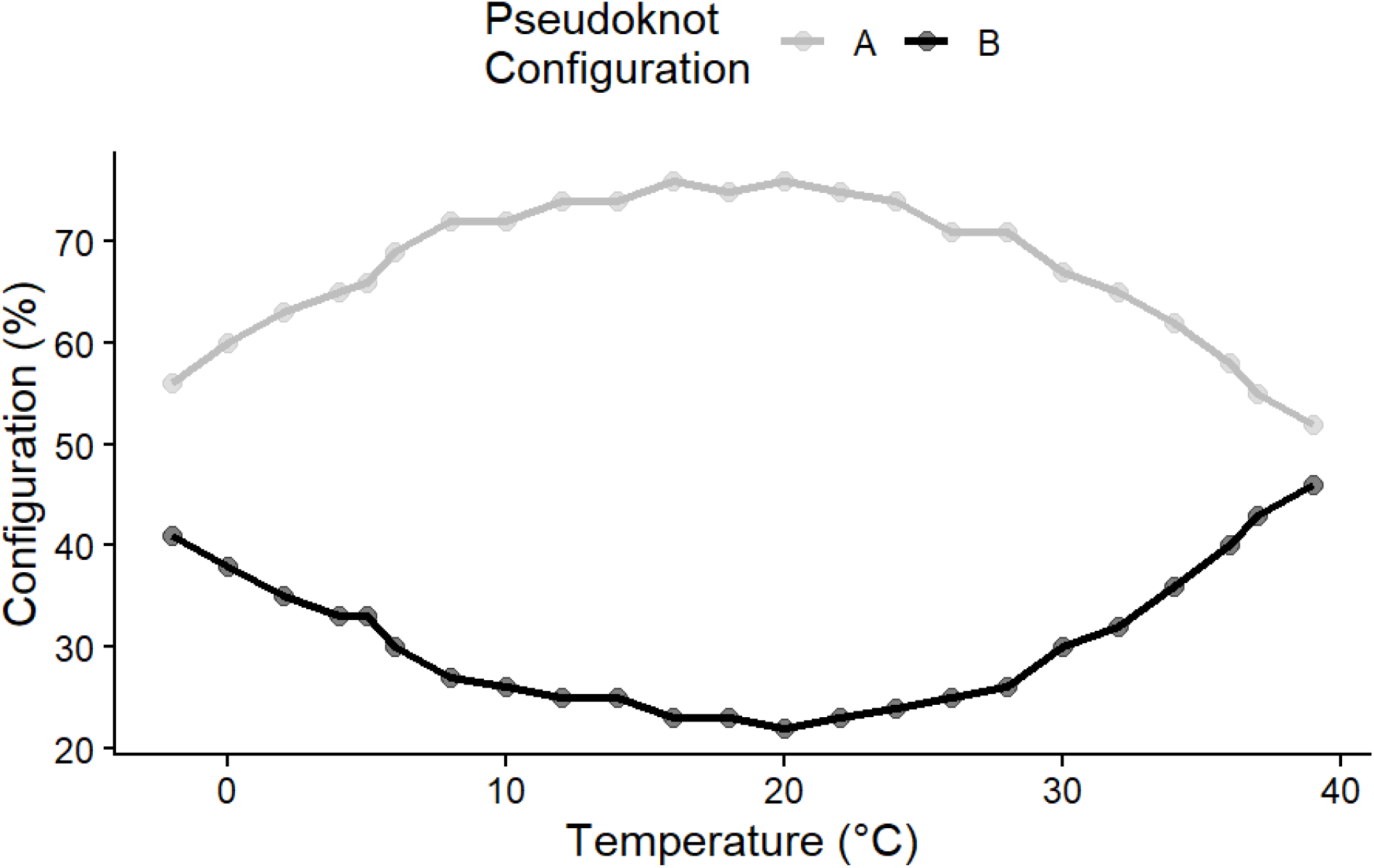
Predicted pseudoknot configuration of Rod 1 region 2 generated using the pKiss “probs” model. Configurations were grouped according to the 5’ initiation site of the primary pseudoknot. In Configuration A, the pseudoknot begins near the 5’ end, whereas in Configuration B, it initiates farther downstream in the sequence. Exemplar of configuration A “.[[[……{{{{{{..((((…))))..]]]..<<<<}}}}}}……>>>>..[[[[….{{{{.]]]]…}}}} “and configuration B “……….[[[[[[[..{{{{{{{(((…….)))]]]]]]]….<<<<….}}}}}}}…>>>>((((….)))).. “.

Rod 1 region 1 had kmer based matches in pre assembled metatranscriptomics data from Alpine Switzerland with fourteen matches between 77.78% and 100% similarity, albeit with abundances too low to assemble (<30), samples are described in Fiore-Donno et al., (2024). MegaBLAST of the alpine datasets revealed that fragmentary reads and reads with up to 2 divergent basepairs were present, but those basepairs were outside of the pseudoknot regions. From the original study in addition to the original find SRX28877766 had 582 hits. Of the sequences with 100% coverage, a 3bp gap outside of the predicted pseudoknot range was identified, and a single high coverage read had only 90% identity to the sequence. Rod 2 region 1 had no kmer based hits, and only one SRX had hits. (SRX28877766), two reads with 100% coverage and identity, which could be chance or contamination. Neither had any homology with the 20 SRA datasets in SRP653358.

## Discussion

Presented are two rod-like structures, Rod 1 has two pseudoknot rich regions which self-complement each other, one of which with conformational temperature shifts that favor low temperatures for pseudoknot folding. Rod 2 has inconsistent conformational results that appear counterintuitive for the source environment. For both structures there is no homology to deltavirus and deltavirus like ribozymes, and the ∼1000 sequences are shorter than known deltaviruses. However, the structural similarities for Rod 1 region 1 from Infernal are significant in the overlapping regions, and the cold temperature conformations bear some structural affinity to deltavirus-like ribozymes and other ribozyme families (nested pseudoknots). While non temperature aware models were inconsistent, the persistence of some pseudoknots across models provides verification for those structures as well as the R-scape conformation for Rod 1 region 1. The megablast hits to insect genomes (34–37 bp) fall below the threshold at which sequence identity alone can establish homology and most likely reflect chance similarity. A transposon origin cannot be formally excluded, but Rod 1 lacks reverse transcriptase coding potential and shares only a 56 bp overlap with the 3839bp non-LTR element Ingi-1_LMi. The non-repetitive character of the remainder of the sequence (assessed by YASS) further argues against a transposon interpretation (Bao & Jurka, 2014).

Based on read presence in at least fourteen samples from a similar ecosystem in the Swiss Alps, presence in a second Finnish sample, the consistency of rod-like structures with the survival of independent RNAs, and pseudoknot confirmation, the putative functional regions are potentially ribozymes or other functional regions analogous to deltavirus ribozymes found in a new grouping of rod-like RNAs. Cross contamination between sites is unlikely as collections were done by different teams, in different countries in different years, with different reagents (Fiore-Donno et al., 2024; Sirja et al., 2025). Likewise, the possibility of a hundred basepair combinations occurring by chance to a read depth of 7 to 15, only in fourteen geographically linked sites and nowhere else in the SRA database is exceedingly unlikely. Rod 2 is less supported by the available data; it appears to exist only in a single location and has inconsistent signalling across models. However, the structural similarity to the Hammerhead ribozyme (type III) is similar to many well documented viroid-like elements whose existence while undocumented in the current environment are unremarkable. (Raza et al., 2025)

Confidence in the exact structure is limited by the possibility that folding predictions currently available are inadequate for analysis of organisms that live exclusively in low temperature conditions where RNA folding patterns differ dramatically (Rissone et al., 2024). However, to the degree that current models can test, the structure persists across biologically relevant temperatures. Modelling outside of default temperatures is well documented for conformational shifts in RNA structure (Eisenhauer et al.,2025). Rod 2 has essentially no confirmation outside of the sample it was found in. Ultimately catalytic activity cannot be confirmed *in silico* and structural similarity to known ribozymes is only suggestive of function.

The nature of several new rod-like RNAs is unknown. Unlike viroids which are well documented in their role as plant pathogens, obelisks and other unclassified rod-like RNAs despite their abundance have no known function and their classification is tentative (Giguère & Perreault; 2017; Urayama, et al., 2025). They could be RNA plasmids associated with organisms, minor pathogens, or relics of RNA life (Done et al., 2026). The latter scenario has been theorized as a potential origin of pre-cellular life and has experimentally driven evolution of RNA polymerase activity (Eigen, 1971; Attwater et al., 2013; Flores et al., 2022). It is notable that captured variation was constrained in Rod 1 region 1, a quasispecies cloud of variation would be expected from a self replicating viroid or viroid like element, however this could be an artifact of low abundance (Sardanyés et al., 2024). Ultimately this finding and findings like this are to be expected with the widespread distribution and diversity of these RNA elements and mapped out pipelines for their detection (Urayama, et al., 2025; Kremer & De Barros, 2024)

## Data Availability

All data used in the study comes from the publicly available SRA database.

## Acknowledgments

The authors wish to thank Dr. Walter Moss for insightful feedback.

## Full Sequences

>Rod2

AGUAGUGUUGUGUUCGGGGAGAAUGGCAGCCCCGCUGAUGAGUCUUAUGCCGAGACGAAACACGAAGACAUAUCGAUACGUAUGGACGUUCUGGUUCCUGUCGUAGCCACCAAGGAGAUAUAUGACUAUGCGCUCUAUGUCUGGCCUUGAGUGUGGAAGACGGCGGUUUCCGGGAGUUCGUACAAGAUCGCGAAGGAGGGUGUAGCAGGGAACAGCGCACUUCGAAGCCCCCGUAAGGUUAACUCGGCCGAUCCCAUACGACUUAAAUGGUCCUUCGUCCCCCAAUGCACAUCGGUCCUCGUAGGUUGAGACCAGGAGAGAAAGAGCCACGACCUAGUCUAAGUGUACAUCCGGCUGUCGAUGAUGAGGUAGUAAAAUAC UUAUGAAUCAUCCGCGUGAGUCCGGGGCACAAUCAAAGGCAGAAGUUGGUACUCUCUUCAAGUAGGCCUAUGGAUGAGGUGAUUCGGUGUGCAGGGUGAGACGUCGGUUACAAGUCAAAAGGAGGUUUCGGCUGGCUGGCCUCAUCAGGGACAACGCGCUACUGUUGCCCUACCUCCCUUACCUUCGUGACCUGUAUCCCCUCCAGUUCCGCAUAUUCCUACCACCAUUCAUAUCACUGUCAUCUGUUGUUCCCUUUCGGCUAUCGGCCGGUUCCAGUUCAUAUGUCCUUUUCGAUAUGCCGCGUGUUUCGUCUUUACGACUAUAGUUGCUAUGCUGCCAGCCCCUCUUACACACGAAGACCUUUUUGAAGCACCAUCACAGCGUGUAGAUAUGAGAUGGCCAAUCAACCGAGGCAGUGUACAGUCGAGUAACGAGGUCGUGGCACCCAGAGAAUUCCGGAGAAGCGCUGAUUGACGAGUUGUUGUUUAAUCCGUCCUUUGCUUCCUCUCGUACUCUAGCUGACCCCCACGCGUGACAAGGGCCUAAAGAAACAACAGGUCAAACGCAAUGUCGCGGGGUAGUCAGUUGUGCCACUAGAUACCGAGCUCCUGAUCGCUGCGGAGGGAUUGAGUAAGUCCUUAAACAACUAGUUUGCAUGUUGACGGGUGUACUUCCACUGGCC

>Rod2region1 AGUAGUGUUGUGUUCGGGGAGAAUGGCAGCCCCGCUGAUGAGUCUUAUGCCGAGACGAAACACGAA

>Rod1 AGCUUACCAGGCAUGGCAAUAACGUGGACGAGAGUCAUACCGGCACCGGCUCAGGUAAGGGAUGUCCUGUAAACAUAUGAUAGCUCUCAGCUUCGGCUGUAUAUUCUCAACAGCCUACUGACAGGGCCCACUACGUGAAAGUGGCUACCAACGGGCCCCCUAAGGACAAAUAUCAGACUUCUCGACUGUCUCACACACUCGCUCUGGAUAACUGCUCUCCGCUUGCCUCACCCACCAUCUCUAGAUUAUCGCAACAACAUCCACUCUUACAAUUGAUCUUCUGAUCGCUGGGGAUCAAUAUCCGAAAGGCCUAUACUACUAAUAGUCGACUGUCUUUCCACCUGCUAAAAGAUGAUGUCCGAGGUUGCAACGCAUGACGAGAUAUCCCGGUUAGAUGGAGCGAUCGGUAAAGGGUAAGUUUCACAGGACAUAAACCCCACAACAACACUGUGUGAGAUCGACAGAAUGUCGUAUUCUGGAAAACACCUCUAUAGCGACGGCUGUUCUACCUUUCCUAUCCAUGGCUUCAGUUGGGUCCUCUUCUCAUAUAGCUGUAAGUAUUCGGAGAGUCUUGUGUAGAUGGUUAGGCUCGUCAACAAAGAGCACACUAUGCCUGGGUCGAAUUGGUUAAAACAACACUUGCCAAGCAGUGGGACGGAUGGAAGCAUCGCUAGAAGGGGAUCGAUGGGGUCGGCAUUAGGAUUACUUCAGAAAAUAGGUGUUGGUUGUGAUGCAGUCUGGAUAUUAUCGUCUAGUCAUCCUCAGGCACCCGCUGGUUUCCAAAGACACAUUAGCGUCUUAUCAGUAAUCCGUUAGGAAGUCUGCUAGAGCGUUCAUCAUAUGUGUUCAGUUCAGAUCUUGUGAGUUAAGGGGUGUUGGUAUGAUGCCUCCACGUGGCGCACCCUGGGUAACGCUAAAUCAUACACUCGUUGGUCAUCGGGCCUUACCGUCUUUUACGCAUUGCGCAAGAGGGGGGACGGGGAUACCACGAGUCGCAUGAGGA

>Rod1region1 UUCGGCUGUAUAUUCUCAACAGCCUACUGACAGGGCCCACUACGUGAAAGUGGCUACCAACGGGCCCCCUAAGGACAAAUAUCAGACUUCUCGACUGUCU

>Rod 1 region 2

UGUGAGUUAAGGGGUGUUGGUAUGAUGCCUCCACGUGGCGCACCCUGGGUAACGCUAA AUCAUACACUCGUUGGUCAUCGGGCCUU

